# Representing Organic Matter Thermodynamics in Biogeochemical Reactions via Substrate-Explicit Modeling

**DOI:** 10.1101/2020.02.27.968669

**Authors:** Hyun-Seob Song, James C. Stegen, Emily B. Graham, Joon-Yong Lee, Vanessa A. Garayburu-Caruso, William C. Nelson, Xingyuan Chen, J. David Moulton, Timothy D. Scheibe

## Abstract

Predictive biogeochemical modeling requires data-model integration that enables explicit representation of the sophisticated roles of microbial processes that transform substrates. Data from high-resolution organic matter (OM) characterization are increasingly available and can serve as a critical resource for this purpose, but their incorporation into biogeochemical models is often prohibited due to an over-simplified description of reaction networks. To fill this gap, we proposed a new concept of biogeochemical modeling—termed *substrate-explicit modeling*—that enables parameterizing OM-specific oxidative degradation pathways and reaction rates based on the thermodynamic properties of OM pools. The resulting kinetic models are characterized by only two parameters regardless of the complexity of OM profiles, which can greatly facilitate the integration with reactive transport models for ecosystem simulations by alleviating the difficulty in parameter identification. For every detected organic molecule in a given sample, our approach provides a systematic way to formulate reaction kinetics from chemical formula, which enables the evaluation of the impact of OM character on biogeochemical processes across conditions. In a case study of two sites with distinct OM thermodynamics, our method not only predicted oxidative degradation to be primarily driven by thermodynamic efficiency of OM consistent with experimental rate measurements, but also revealed previously unknown critically important aspects of biogeochemical reactions, including their condition-specific response to carbon and/or oxygen limitations. Lastly, we showed that the proposed substrate-explicit modeling approach can be synergistically combined with enzyme-explicit approach to provide improved predictions. This result led us to present *integrative biogeochemical modeling* as a unifying framework that can ideally describe the dynamic interplay among microbes, enzymes, and substrates to address advanced questions and hypotheses in future studies. Altogether, the new modeling concept we propose in this work provides a foundational platform for unprecedented predictions of biogeochemical and ecosystem dynamics through enhanced integration with diverse experimental data and extant modeling approaches.

## INTRODUCTION

Organic matter (OM) is a key determinant of global biogeochemistry and exerts far-reaching impacts on regional and global ecosystem health. Degradation of complex OM is primarily driven by microbial and enzymatic activities around environmental substrates (Moorhead et al., 2013;Manzoni et al., 2016;Paul, 2016;Varjani, 2017). Therefore, a proper representation of the dynamic interplay among microbes, enzymes, and substrates is key for reliable prediction of OM cycling and ecosystem functioning. Despite an increasing ability to generate high-resolution data for each of these components, understanding of fundamental processes that govern their interplay is limited and, consequently, is poorly represented in models (Fatichi et al., 2019). To date, no current-generation modeling framework has been available to interpret and utilize increasingly available high-resolution metabolite data for predicting biogeochemical cycles at the ecosystem level.

Earlier biogeochemical models provide a lumped description of microbial and enzymatic activities, as well as chemical entities (Schimel and Schaeffer, 2012;Blankinship et al., 2018). This reductionist approach significantly limits the predictive ability of models. For the past decade, growing recognition of the significance of microbial activities on biogeochemistry has led to the development of microbial-explicit models (MXMs) (Todd-Brown et al., 2012;Wieder et al., 2015;Allison, 2017;Sulman et al., 2018), which include DEMENT (Allison, 2012), CORPSE (Sulman et al., 2014), MIMICS (Wieder et al., 2014), MEND (Wang et al., 2015), RESOM (Tang and Riley, 2015), and other functional guild-based models (Hood et al., 2006;Jin and Roden, 2011;Bouskill et al., 2012). With or without coupled consideration of enzymatic processes, these models account for microbial physiology and interactions to provide a deeper understanding of microbially-mediated OM decomposition.

Enzyme-based or enzyme-explicit models (EXMs) are a complementary approach that collectively describes biogeochemical functions performed by a microbial community with a focus on extracellular enzymes, alleviating the difficulty in identifying functional traits of individual organisms or their groups (Moorhead et al., 2013;Song et al., 2014;Song and Liu, 2015;Song et al., 2017). In contrast with such increasing details considered in MXM and EXM, over-simplified descriptions of substrate pools remain as a serious *bottleneck* in building predictive biogeochemical models because further elaboration of microbial and enzyme activities becomes difficult without expanded consideration of complex OM chemistry that influences the specific metabolic pathways employed by microorganisms.

Extension of biogeochemical models to include detailed OM chemistry is critically important to pushing the boundaries of environmental science forward. For example, a present paradigm in environmental science views aerobic respiration rates as being primarily determined by kinetics (i.e., organic carbon (OC) and oxygen concentrations). However, recent field and laboratory studies suggest that OM thermodynamics can be a main driver of aerobic respiration (Graham et al., 2017;Graham et al., 2018;Stegen et al., 2018b;Garayburu-Caruso et al., 2020). Conventional lumped biogeochemical models are not completely effective for addressing this issue because the thermodynamic properties of substrates are a function of chemical composition of compounds constituting OM pools. This implies that identification of underlying key processes that drive aerobic respiration requires advanced biogeochemical models that can properly reflect all relevant kinetics and thermodynamics in representing actual oxidation rates.

Towards filling this gap, we propose a new modeling concept drawn from thermodynamic theory that can explicitly account for the chemical composition of individual molecules in OM pools when generating biogeochemical rate estimates, therefore having the potential to significantly advance predictive capabilities of current-generation ecosystem models. Due to the ability of our approach to directly incorporate OM chemistry for every compound, we termed it *substrate-explicit modeling* (SXM), as opposed to MXM and EXM. SXM features flexibility in input data types, enabling incorporation of an unlimited number of compounds detected from various state-of-the-art instrumentation including GC-MS, LC-MS/MS, HPLC-MS, NMR, Orbitrap MS, and Fourier transform ion cyclotron resonance (FTICR-MS). Our modeling framework therefore overcomes a central challenging in using such information—how to condense the massive amount of data produced by these technologies into variables that are both useful and computationally feasible in predicting biogeochemical dynamics and function.

By combining a suite of previously developed thermodynamic theories (McCarty, 2007;Kleerebezem and Van Loosdrecht, 2010;LaRowe and Van Cappellen, 2011;Desmond-Le Quemener and Bouchez, 2014), we developed a systematic procedure to convert chemical formulae of all organic compounds detected in an environment, regardless of the number of compounds or measurement technique, into two rate parameters that integrate seamlessly into a variety of modeling constructs. We evaluated the effectiveness of our approach through the analysis of OM characterized via FTICR-MS data obtained from two biogeochemically distinct sites. Model outputs were compared to experimental work on the same samples by Graham et al. (2017; 2018), who showed that aerobic respiration varied between across sites by several-fold range. We chose to model two representative OM profiles from the samples with high (high activity, HA) and low (low activity, LA) rates of aerobic respiration to test model predictions against the broadest range of biogeochemical activity in the dataset.

Consistent with the findings from Graham et al. (2017; 2018), comparative analyses of the two SXMs constructed for HA and LA zones showed that OM pools in HA zones were thermodynamically more favorable than those in LA zones and that this difference of OM thermodynamic properties was associated with elevated respiration rates in the HA zone. Further comparison of predicted reaction rates and experimentally determined aerobic respiration enabled reversely inferring field conditions where data were collected. Lastly, we showed that SXMs have a flexible structure to be synergistically integrated with other existing frameworks (such as EXMs and/or MXMs) towards more comprehensive predictive biogeochemical modeling. Through coupling with reactive transport models, this hybridized or integrative biogeochemical modeling (IBM) is expected to significantly expand our ability to predict complex ecosystem functions.

## METHODS

Our development of SXMs from chemical formula of OC go through multiple-step procedures as illustrated in (Figure 1a), which are classified into two major parts: (1) derivation of stoichiometric equations for catabolic, anabolic, and metabolic reactions by combining a set of standard thermodynamic analyses (McCarty, 2007;Kleerebezem and Van Loosdrecht, 2010;LaRowe and Van Cappellen, 2011); and (2) formulation of kinetic equations for the final oxidative degradation reaction of OC using a relatively recent thermodynamic theory for microbial growth (Desmond-Le Quemener and Bouchez, 2014).

**Figure 1.**
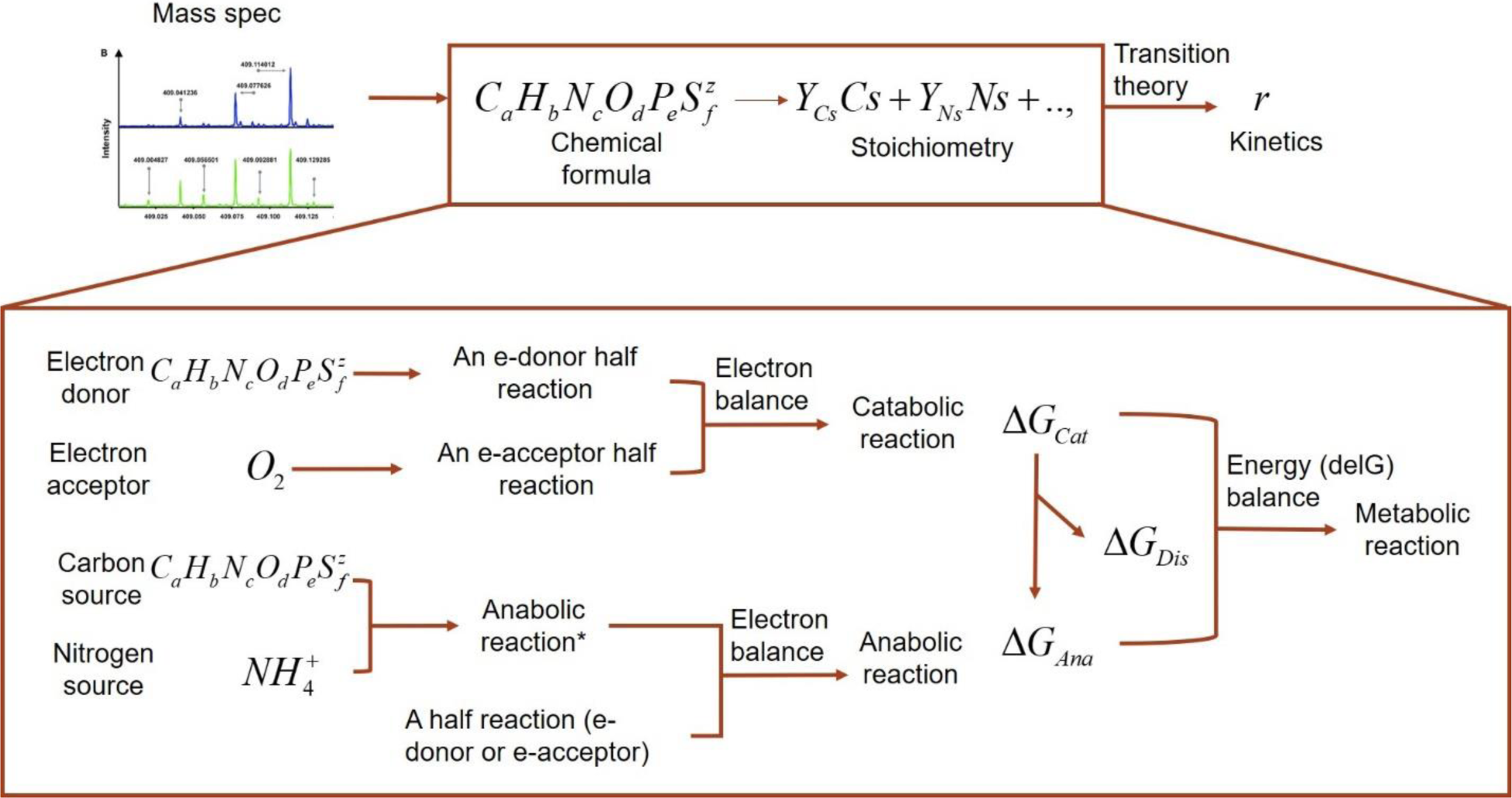
A schematic illustrating the flows of building substrate-explicit models from chemical formulae of OC to stoichiometry and kinetics of oxidative respiration. The zoomed-in box below shows the bottom-up derivation of biogeochemical reactions for each OC.

### Stoichiometric representation of oxidative respiration reaction

Stoichiometric equations provide quantitative relationships among the reactants and products involved in given reactions. We derived a stoichiometric equation for oxidative degradation of OC by accounting for catabolism (i.e., all processes for obtaining energy through substrate oxidation or other means) and anabolism (i.e., synthesis of biomass using the energy provided from catabolism) (Figure 1b). Combination of catabolic and anabolic reactions through energy coupling leads to metabolic reactions. Catabolic reactions (and anabolic reactions as well in many cases) are in turn combinations of a pair of redox half reactions, i.e., for an electron donor (Ed) and an electron acceptor (Ea). Therefore, oxidative degradation of OC described here as a metabolic reaction can be systematically derived in a bottom-up fashion. We provide step-by-step procedures of developing stoichiometric equations following the standard approaches outlined in the literature (Kleerebezem and Van Loosdrecht, 2010;Rittmann and McCarty, 2012). For a general representation of all associated reactions, we use the following convention for stoichiometric reaction (*R*):

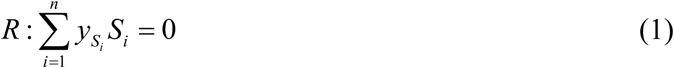

where *S*_*i*_ denotes chemical species *i*, 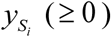 is the stoichiometric coefficient of *S*_*i*_, which is set to be negative for reactants and positive for products. For a given array of *S*_*i*_ ‘s, the stoichiometric equation in Eq. (1) can be conveniently represented as a column vector:

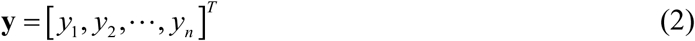

where the superscript *T* denotes the vector transpose. In the context of OC oxidation reaction, we specified 10 chemical species (*C*_*i*_ ‘s) to define *R* (Table 1).

**Table 1:** Key chemical species that participate in oxidative degradation of OC and their standard Gibbs free energy of formation at pH=0.

| $i$ | $S_i$ | $G^{f0}$ [kJ/mol]** | Source for $G^{f0}$ value |
| --- | --- | --- | --- |
| 1 | $OC^*$ | Unknown | |
| 2 | $H_2O$ | -237.2 | (Alberty, 2006) |
| 3 | $HCO_3^-$ | -586.9 | (Alberty, 2006) |
| 4 | $NH_4^+$ | -79.4 | (Kleerebezem and Van Loosdrecht, 2010) |
| 5 | $HPO_4^{2-}$ | -1096.1 | (Alberty, 2006) |
| 6 | $HS^-$ | 12.1 | (Kleerebezem and Van Loosdrecht, 2010) |
| 7 | $H^+$ | 0 | (Kleerebezem and Van Loosdrecht, 2010) |
| 8 | $e^-$ | 0 | (Kleerebezem and Van Loosdrecht, 2010) |
| 9 | $O_2$ | 16.4 | (Alberty, 2006) |
| 10 | $Biom^{**}$ | -67 | (Kleerebezem and Van Loosdrecht, 2010) |

\* $OC$ = organic carbon; \*\* biomass: $CH_{1.8}N_{0.2}O_{0.5}$ (Stephanopoulos et al., 1998; Kleerebezem and Van Loosdrecht, 2010); \*\*\* Standard formation values for the Gibbs energy

#### Catabolic reaction

Catabolic reaction is obtained by combining Ed (i.e., OC) and Ea (i.e., O_2_) half reactions (denoted by *R*^*D*^ and *R*^*A*^). Following (LaRowe and Van Cappellen, 2011), we write *R*^*D*^ as follows:

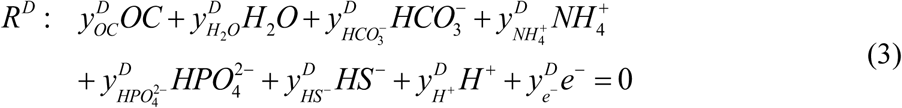

The chemical formula for OC is represented as follows:

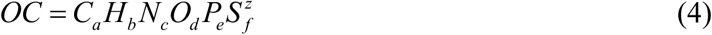

where the subscripts *a, b, c, d, e*, and *f* denote the elemental composition of OC in terms of C, H, N, O, P, and S, and the superscript *z* represents the net charge of OC. Stoichiometric coefficients in Eq. (3) are functions of elemental composition of OC (i.e., *a, b, c, d, e, f*, and *z*) (LaRowe and Van Cappellen, 2011). The Ea half reaction *R*^*A*^ is given in a simple form as follows (Rittmann and McCarty, 2012):

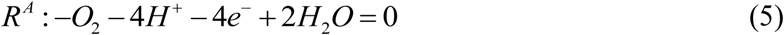

These reactions can be represented in terms of the following two vectors of stoichiometric coefficients, i.e.,

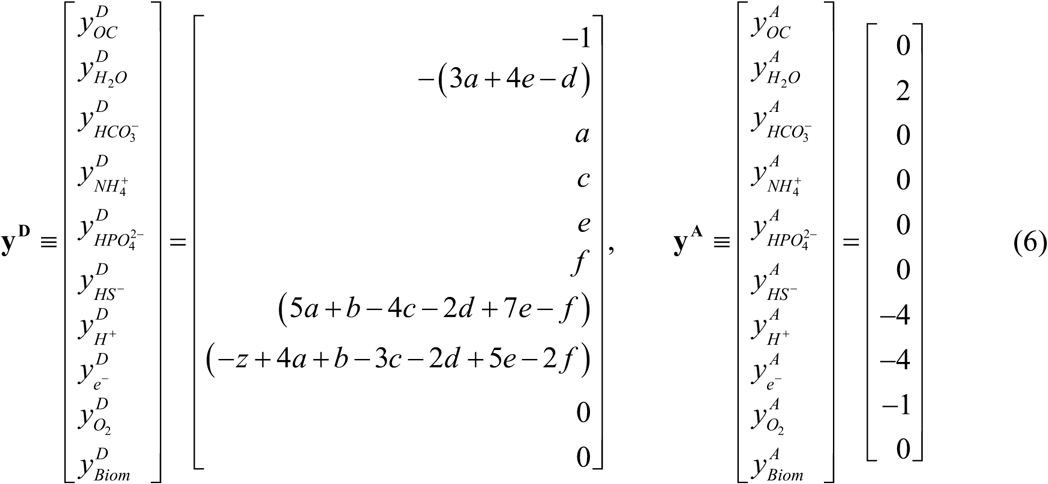

The overall stoichiometry of the complete catabolic reaction equation (*R*^*Cat*^) is obtained by combining the Ed and Ea half reactions:

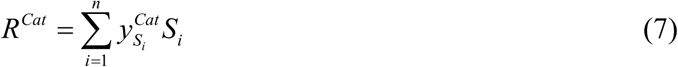

where the stoichiometric coefficient vector is determined as follows such that the net electron production or consumption through *R*^*Cat*^ is zero, i.e.,

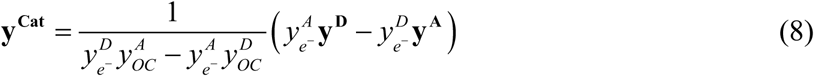

#### Anabolic reaction

In anabolism, the substrates (carbon source and nitrogen source) are reduced to biomass. When the carbon source is the same as the electron donor (i.e., *OC*) (which is true for heterotrophic bacteria) and the nitrogen source is ammonium, the anabolic reaction equation (*R*^*An*\*^) can be written as follows:

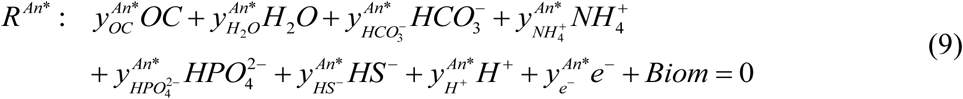

Similar to previous cases, the vector of stoichiometric coefficients in *R*^*An*\*^ (denoted by **y**^**An\***^) can also be represented as a function of composition of OC to meet elemental mass balances, i.e.,

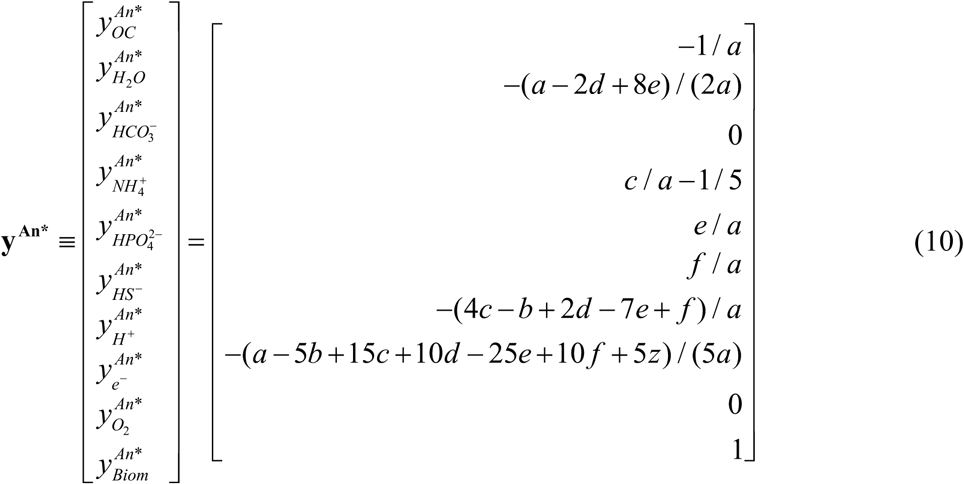

Note that 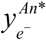 can be zero, negative or positive depending on the difference of oxidation states between the biomass and substrates (denoted by *γ*_*Biom*_ and *γ*_*Sub*_). If *γ*_*Biom*_ < *γ*_*Sub*_ (or 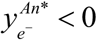), it implies electrons are required to convert substrates to biomass; if *γ*_*Biom*_ > *γ*_*Sub*_ (or 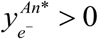), the conversion of substrates to biomass produces electrons, requiring an electron acceptor. In combination with electron donor or acceptor equation defined in Eqs. (3), (5), and (6), the anabolic reaction equation (*R*^*An*^) is finally represented without the electron term as follows:

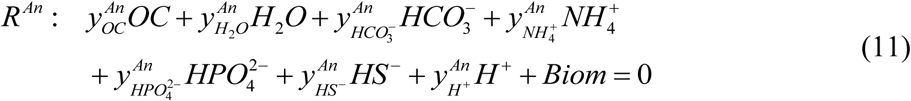

where the vector of stoichiometric coefficients is given as follows:

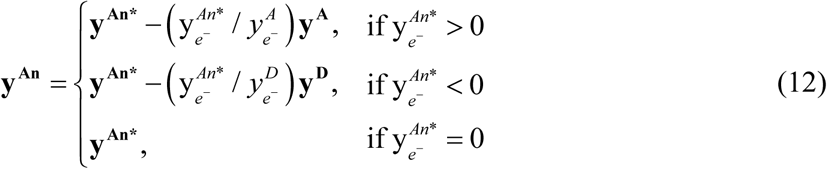

#### Metabolic reaction

Finally, a metabolic reaction equation is obtained by accounting for the energetic coupling between catabolism and anabolism, i.e., the balance between the energy production through substrate degradation and the energy consumption for cell synthesis (i.e., biomass production). Two approaches commonly considered for this purpose include the dissipation method (Heijnen et al., 1992;Heijnen and Vandijken, 1993) and the thermodynamic electron equivalents model (TEEM) (McCarty, 2007). We used the TEEM in this work because the dissipation method is applicable for up to six-carbon OCs, while most of the compounds profiled in our samples are more complex than those. The TEEM considers the energy provision of catabolic reaction to meet the energy spent in the following two steps of energy conversion in anabolism: (1) the carbon source to biomass building blocks (Δ*G*^*Block*^), and (2) the conversion of biomass building blocks to biomass (Δ*G*^*Syn*^). The original formulation includes the energy conversion from the nitrogen source to ammonium, which is neglected in our case where ammonium is taken as the nitrogen source. The energy balance is then represented using the following equation:

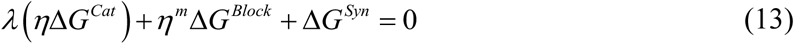

where Δ*G*^*Cat*^ is the Gibbs free energy change in *R*^*Cat*^, *λ* is a parameter that couples catabolism and anabolism, and *η* is an energy transfer efficiency parameter. The exponent *m* accounts for energy transfer efficiency depending on whether energy is generated or consumed in the process of converting the carbon source to biomass building blocks by setting it to +1 (if Δ*G*^*Block*^ < 0) or −1 (if Δ*G*^*Block*^ > 0). Following (Kleerebezem and Van Loosdrecht, 2010), we chose Δ*G*^*Syn*^ = 200 *kJ* / (*mol Biom*), and *η* = 0.43; assumed the composition of biomass building block to be the same as *Biom* (i.e., *CH*_1.8_ *N*_0.2_*O*_0.5_).

The stoichiometric coefficient vector for metabolic reaction (**y**) is obtained by coupling the catabolic and anabolic reactions using *λ*, i.e.,

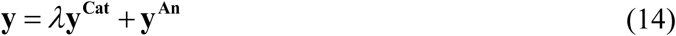

where the parameter *λ* implies how many times the catabolic reaction needs to run in order to provide the energy required for the synthesis of a unit C-mole of biomass. Determination of *λ* from Eq. (13) requires the calculation of the Gibbs free energy changes such as Δ*G*^*cat*^ and Δ*G*^*Block*^, the latter of which is the same as Δ*G* ^*An*^ in our formulation (where the elemental composition of biomass building block was treated as the same as *Biom*). In the following section, we describe how to calculate the Gibbs free energy changes in a general context.

### Gibbs free energy change

The Gibbs free energy change of a chemical reaction (Δ*G*) can be calculated based on the formation energy of all participating compounds 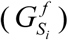 and their stoichiometry 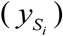. For dilute aqueous systems, Gibbs free energy is often calculated at the standard condition (as denoted by the superscript ‘0’ in the following equation), i.e.,

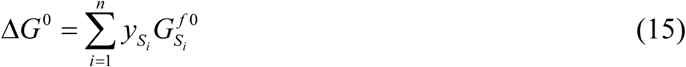

The standard state of dissolved compounds in aqueous systems is defined at 1 mol/L and zero ionic strength. However, the hydrogen ion (*H*^+^) at that standard concentration implies pH = 0, which does not properly represent the condition where actual biochemical reactions take place. Thus, we instead used the Gibbs free energy for the biochemical standard state at the condition where all compounds are at 1 mol/L but [*H*^+^] = 10 ^−7^ mol/L (i.e., pH = 7). This gives the biochemical standard Gibbs free energy change (denoted by Δ*G*^0′^) as follows:

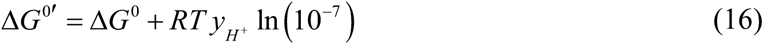

where *R* is the gas constant (= 0.008314 kJ/(K·mol)), *T* is temperature (=298.15 K at the standard condition). It is generally regarded that Δ*G*^0′^ is sufficiently accurate for thermodynamic analysis of redox reactions in biochemical systems (Kleerebezem and Van Loosdrecht, 2010). Hereafter, we drop the superscripts 0′ to denote thermodynamic functions at pH=7, while we still keep the superscript 0 to denote their values at pH=0.

For all compounds considered in this work except OC, the values of 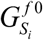 in Eq. (15) are readily obtainable from public databases (Flamholz et al., 2012) or the literature (see Table 1) (Alberty, 2006;Kleerebezem and Van Loosdrecht, 2010). Theoretical estimation of the standard formation energy of OC 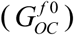 using the group contribution theory is infeasible because the structural information of compounds is not available in FTICR-MS. Therefore, we estimated 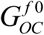 in the following way.

First, we estimated the standard Gibbs free energy change for an electron donor half reaction (Δ*G*^*D*0^) using the formula from (LaRowe and Van Cappellen, 2011). They provided a linear relationship between the nominal oxidation state of carbon (NOSC, denoted by *γ* here) and the Gibbs energies for the oxidation half reactions of OC represented on a C-mole basis (Δ*G*^*Cox*0^), i.e.,

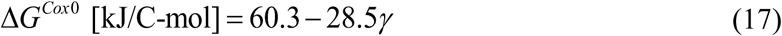

where *γ* is obtained from the exchanged electron moles in the half reaction, and the number of carbon in OC:

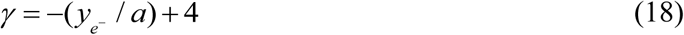

As Δ*G*^*Cox*0^ represents the Gibbs free energy per one C-mole, Δ*G*^*D*0^ is obtained simply by multiplying the number of carbon in OC, i.e.,

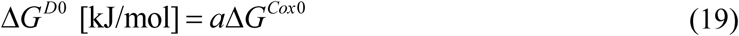

Then, the substation of Δ*G*^*D*0^ in the above into Eq. (15) leads to

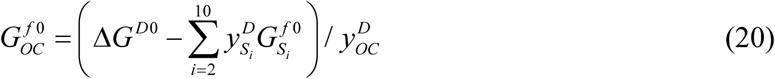

where 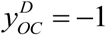 as shown in Eq. (6). Once 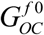 is known from Eq. (20), it is straightforward to calculate the standard (at pH=0) and biochemical standard (at pH=7) Gibbs free energy changes for any given reactions.

### Reaction kinetics

The foregoing sections show how to systematically convert chemical formula of OC to stoichiometric equations for oxidative OC degradation reactions including catabolic, anabolic, and metabolic reactions. As a next step, we extend it to formulate kinetics using the microbial growth thermodynamic theory developed by Desmond-Le Quemener and Bouchez (2014). They considered microbial growth through the following two steps: (1) reversible transition of a microbe (*X*) to an activated state (*X* ^‡^) and (2) irreversible cell division from the activated cell to two daughter cells, i.e.,

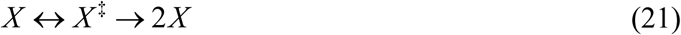

Similar to the classical transition state theory in chemical reaction (Truhlar et al., 1996), the reversible reaction in the first step is assumed to be faster than the second step so that they are in equilibrium. During the first step, each microbe harvests chemical energy from environment. In order to trigger cell division, the energy acquisition from environment should exceed a certain threshold level (*E*^‡^, activation energy). The activation energy is nothing but the summation of anabolic energy requirement and energy dissipation, i.e.,

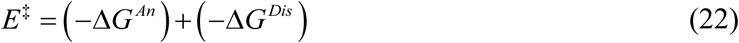

The total usable energy for growth depends on two factors: (1) the energy generation through catabolism (Δ*G*^*Cat*^), and (2) the availability of energy sources (i.e., substrates) in environment. The second factor depends on substrate concentration ([*S*]) and the volume that a microbe can access for harvesting energy (*V*_*h*_, harvest volume). Using statistical analysis, Desmond-Le Quemener and Bouchez showed that the equation for microbial growth rate (*μ*) can be formulated as follows:

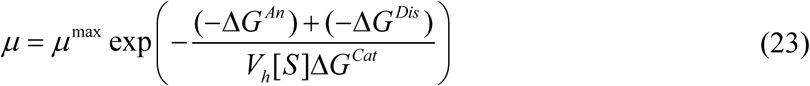

where *μ*^max^ is the maximal specific growth rate. The above equation can be rewritten in terms of *λ* defined in Eq. (13), i.e.,

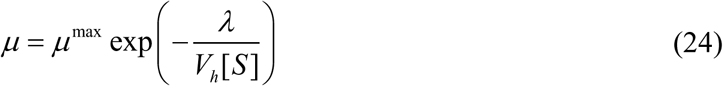

In our current formulation where all stoichiometric equations were derived for a unit C-mole of biomass, the negative value of the parameter *λ* is equal to the stoichiometry coefficient of OC. For the *i*^*th*^ OC (i.e., *OC*_*i*_), this leads to the following form of microbial growth rate:

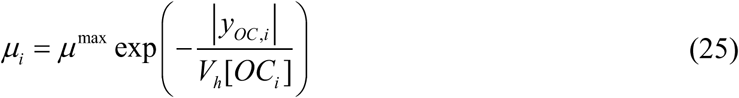

where *μ*_*i*_ is the microbial growth rate on *OC*_*i*_, *y*_*OC,i*_ is the stoichiometric coefficient of *OC*_*i*_ in the metabolic reaction, and |*y*_*OC,i*_ | denotes the absolution value of *y*_*OC, i*_. In the equation above, we assumed that the maximal growth rate remains the same across

Desmond-Le Quemener and Bouchez also provided the extension of the formulation to multiple substrates. In the case of oxidative degration of OC, the final growth equation becomes

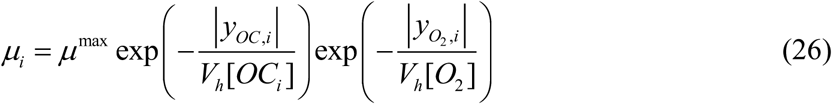

where 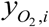 is the stoichiometric coefficient of O_2_ in the metabolic reaction associated with oxidative degradation of *OC*_*i*_. The above equation shows the case when both C and O_2_ are limited. If only C or O2 is limited, Eq. (26) is reduced to

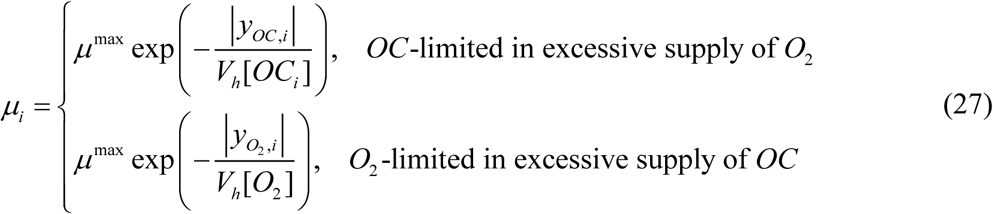

Consumption and production rates of other chemicals (such as *OC, C, O*_2_, and 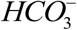) can be obtained simply by multiplying stoichiometric coefficients and *μ*_*i*_ in Eq. (25), i.e.,

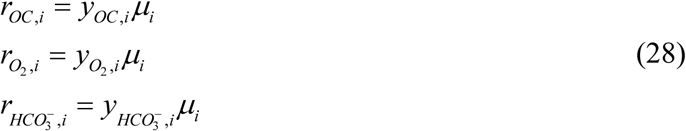

Finally, it is straightforward to set up dynamic mass balances. For example, in a homogeneous batch configuration, dynamic SXMs for key metabolites that are associated with oxidative respiration of *OC*_*i*_ can be written as follows:

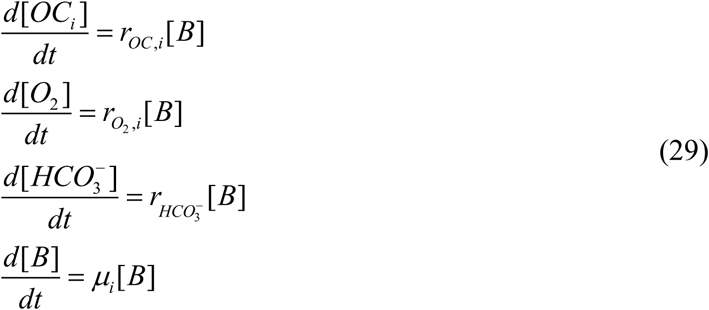

### Integration with MXM and EXM

The SXM derived in Eq. (29) can be combined with other complementary approaches. Integration of SXM with MXM requires explicit consideration of distinct microbial groups, which leads to

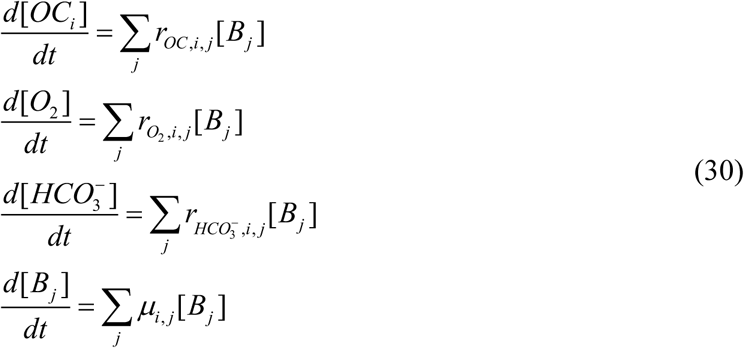

where the subscript *j* denotes the contribution of the *j*^*th*^ microbial group to the production and consumption of metabolites, and *μ*_*i, j*_ is the growth rate of the *j*^*th*^ microbial group on *OC*_*i*_, i.e.,

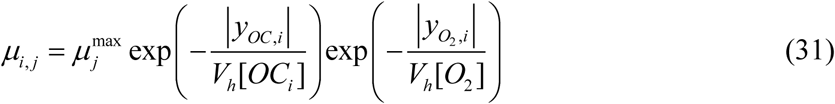

Note that this formulation accounts for the difference in growth rate among microbial groups. Production and consumption rates of metabolites are accordingly dependent on microbial groups, i.e.,

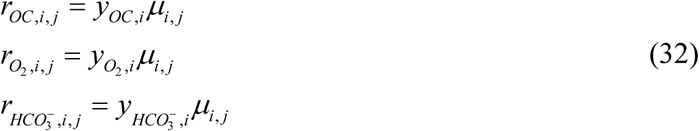

For the integration of SXM with EXM, we introduce enzyme concentrations as additional variables. Dynamic mass balances in Eq. (29) remain the same, but we consider individual reactions to be catalyzed by distinct enzymes, i.e.,

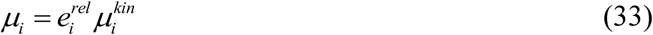

where the reaction rate (*μ*_*i*_) is composed of two parts: enzymatic regulation 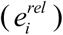 and kinetic conversion 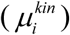, and 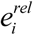 denotes relative level of enzyme, i.e.,

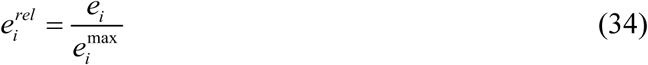

Enzyme concentrations and their maximal values can be determined from the following equation:

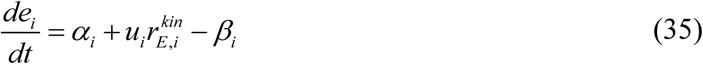

where three terms on the right hand side of the above equation respectively denote constitutive enzyme synthesis rate, inductive enzyme synthesis rate, and enzyme degradation rate, the variable *u*_*i*_ is the cybernetic variable that controls inductive enzyme synthesis, and 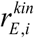 is the kinetic rate of enzyme synthesis. The cybernetic variable *u*_*i*_ can be determined from the Matching Law (Ramkrishna and Song, 2012;Ramkrishna and Song, 2018). Instead of solving separate enzyme equations, we considered a simplified cybernetic model (Song et al., 2018) where 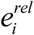 is approximated by *u*_*i*_ (Song and Ramkrishna, 2010;2011), which is in turn determined based on 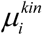. Then, Eq. (33) becomes

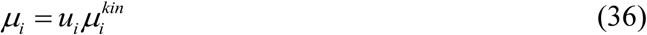

where

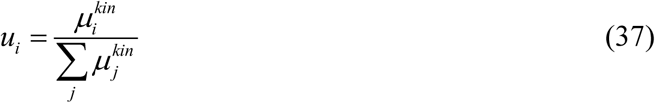

### Field samples

To demonstrate our modeling concept described above, we used the datasets generated in a previous work, described in detail in Graham and colleagues (Graham et al., 2017;Graham et al., 2018). Briefly, sediment profiles (0–60 cm) were collected along two shoreline transects perpendicular to the Columbia River within the Hanford Site 300 Area in eastern Washington State (Graham, Crump, et al., 2016; Graham et al., 2017; Slater et al., 2010; Zachara et al., 2013): one with riparian vegetation (HA zone) and the other without (LA zone). In each transect, profiles were collected from three locations: upper, mid, and lower banks, and sectioned into 10 cm vertical intervals. FTICR-MS was used to characterize C chemistry in each sample, and Raz reduction assay was performed as a proxy of the rate of aerobic respiration per sample. For each peak detected in FTICR-MS spectra, we assigned a chemical formula using the following steps: (1) transformation of raw spectra to m/z (i.e., mass divided by charge number) values using BrukerDaltonik software; (2) chemical formula assignment using in house software following the Compound Identification Algorithm (Kujawinski and Behn, 2006;Minor et al., 2012;Tfaily et al., 2017). In the case that one m/z value can be matching with multiple chemical formulae, consistent assignment was made based on a set of prescribed rules, including the preference of the formula with the lowest error and with the lowest number of heteroatoms and the requiring the presence of at least four oxygen atoms for the assignment of one phosphorus atom. Peaks not satisfying these criteria were not assigned chemical formulae.

For detailed analysis and comparison of two sites, we chose one sectioned sample from the LA and HA profiles: (1) Upper bank at unvegated sited (N1), 40–50 cm depth interval (sample N1-40-50) and (2) Upper bank at vegetated site (S1), 0–10 cm depth interval (sample S1-00-10). These two samples respectively represent the lowest and highest activity in aerobic respiration except an outlier (see **Results: Comparison of low-activity and high-activity samples**).

## RESULTS

### Evaluation of thermodynamic parameters as an indicator of oxidative respiration

Previous studies have shown that aerobic respiration rates from HA samples were higher than LA samples and provided the hypothesis that thermodynamic favorability of OM might be a key factor in regulating respiration rate (Graham et al., 2017;Graham et al., 2018). While the original papers provided datasets that convincingly support this hypothesis, the question of what thermodynamic parameters (among many introduced in Methods) can serve as reliable indicators of aerobic respiration remains unknown. We therefore chose multiple thermodynamic parameters and examined to what extent individual parameters are correlated with aerobic respiration. Three key parameters include Gibbs free energy changes for electron donor half reactions (Δ*G*^*D*^), Gibbs free energy changes for catabolic reactions (Δ*G*^*Cat*^), and the parameter quantifying the energy coupling between catabolic and anabolic reactions (λ). The first two parameters (Δ*G*^*D*^ and Δ*G*^*Cat*^) evaluate the thermodynamic character of OM with a focus on the energy *generation* (through a half or complete catabolic reaction, respectively). By contrast, the parameter λ quantifies the same based on the energy *balance* (i.e., the total energy generation to meet the demand for the synthesis of a unit C-mole of biomass). Therefore, all of these three parameters measure thermodynamic *inefficiency* of OM (in the sense that lower values of parameters imply higher thermodynamic efficiencies of OM), but at different levels, i.e., half/complete catabolic reactions for Δ*G*^*D*^ and Δ*G*^*Cat*^, and entire metabolic reaction for λ. We examined their correlations with experimentally determined aerobic respiration at pH=0 and pH=7 (Figure 2). We also examined whether the thermodynamic favorability of compound can be better characteized on a C-mole or OM-mole basis (Supplementary Figure S1).

**Figure 2.**
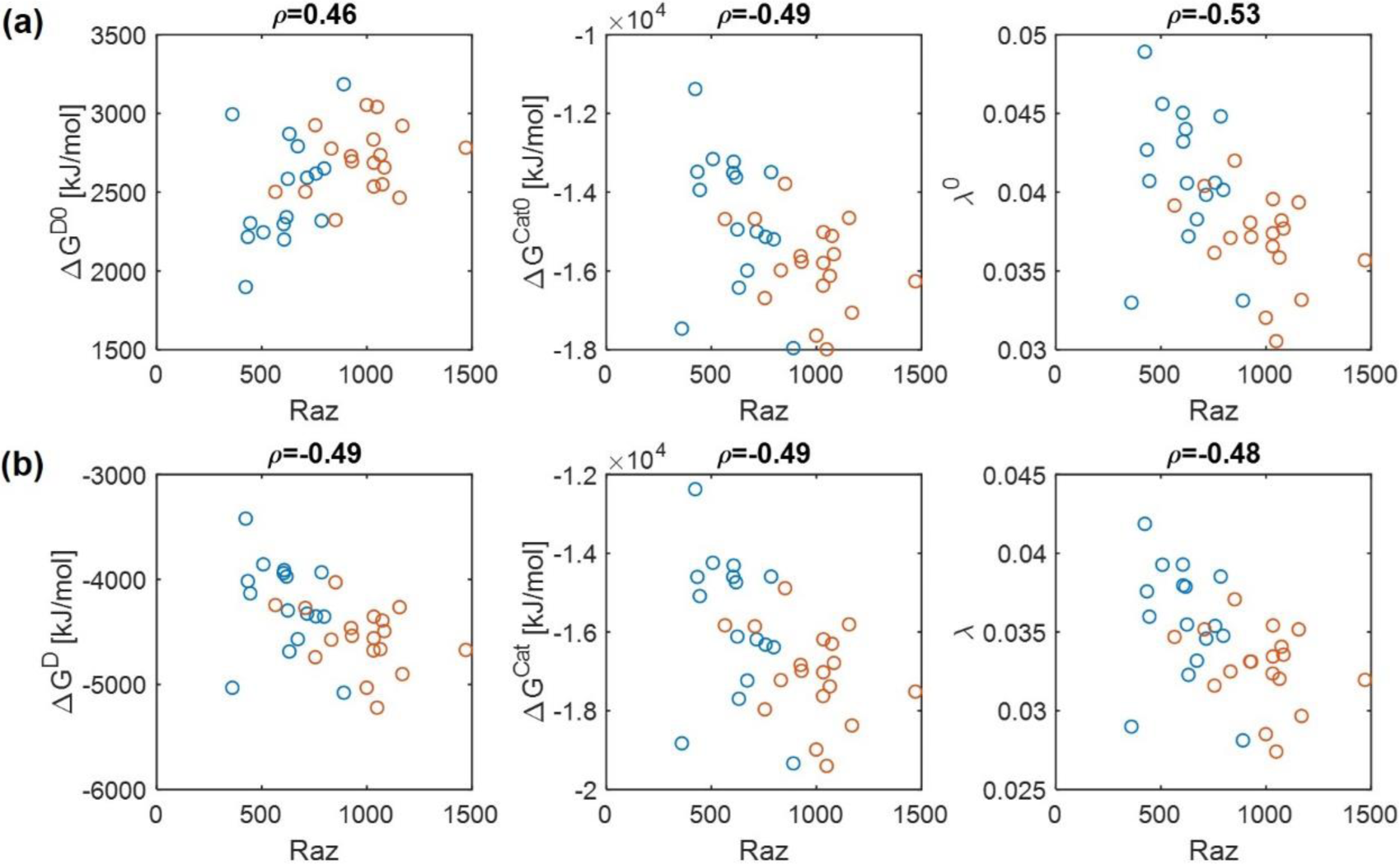
Pearson correlations of aerobic respiration with three key thermodynamic functions: Gibbs free energy changes for an electron donor half reaction (Δ*G*^*D*^) and catalytic reaction (Δ*G*^*Cat*^), and the energy coupling parameter (*λ*): (a) pH=0, standard condition assuming [*H* ^+^] = 1*M* (denoted by the superscript 0) and (b) pH=7, biochemical standard condition assuming[*H* ^+^] = 10^−7^ *M*.

Regardless of pH values, Δ*G*^*Cat*^ and λ showed negative correlations with aerobic respiration, which implies that thermodynamically favorable OM leads to higher respiration rates (the middle and right panels in Figures 2a and 2b). By contrast, the results for Δ*G*^*D*^ were pH-dependent, i.e., Δ*G*^*D*^ showed a positive correlation with aerobic respiration at pH=0 (the left panel of Figure 2a) (indicating that Δ*G*^*D*^ is not a good estimator of aerobic respiration under this condition), while it showed a negative correlation at pH=7 (the left panel of Figure 2b).

As C-mole, the parameter λ still showed negative correlations with aerobic respiration at both pH values. Δ*G*^*D*^ and Δ*G*^*Cat*^ showed weak relationships with aerobic respiration with correlation coefficients between −0.1–0.1, except for Δ*G*^*D*^ at pH=7 where the correlation coefficient was positive.

Together, these results identify the parameter λ as the most robust indicator of aerobic respiration, which shows consistent correlations across all conditions considered above. The results also indicate that the usefulness of the other thermodynamic parameters (Δ*G*^*D*^ and Δ*G*^*Cat*^) as a respiration indicator is relatively limited.

### Model validation by comparing predicted reaction rates with aerobic respiration

Comparison of predicted reaction rates with experimentally determined aerobic respiration provides a means to validate the model. The experiment of Raz reduction assay in Graham and colleagues (Graham et al., 2017;Graham et al., 2018) was designed as proxy measurement for the rate of oxygen consumption. If properly formulated, our model is expected to predict oxygen consumption rates that are positively correlated with experimental estimation. As reaction rates are functions of substrate concentrations, we performed this comparison under three different limiting conditions: (1) C, (2) O_2_, and (3) both C&O_2_ limitation. For each, we differentiated the level of limitation to be severe vs. moderate.

For simplicity, we set *μ*^max^ = 1 in Eqs. (26) and (27) because this value does not affect the correlation with aerobic respiration, while other parameters and variables such as *V*_*h*_, [*OC*], and [*O*_2_] are unknown. To implement different levels of substrate limitation, we set *V*_*h*_ [*OC*] and/or *V*_*h*_ [*O*_2_] to be 1 (moderate limitation) (Figure 3) and 0.2 (severe limitation), respectively (Figure 4) in the following four reaction rates: biomass production (i.e., growth) rate (*μ*), C consumption rate (*r*_*C*_ ≡ (# of C) × *r*_*OC*_), O_2_ consumption rate 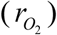, and inorganic carbon production rate 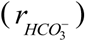 as defined in Eqs. (26) to (28).

**Figure 3.**
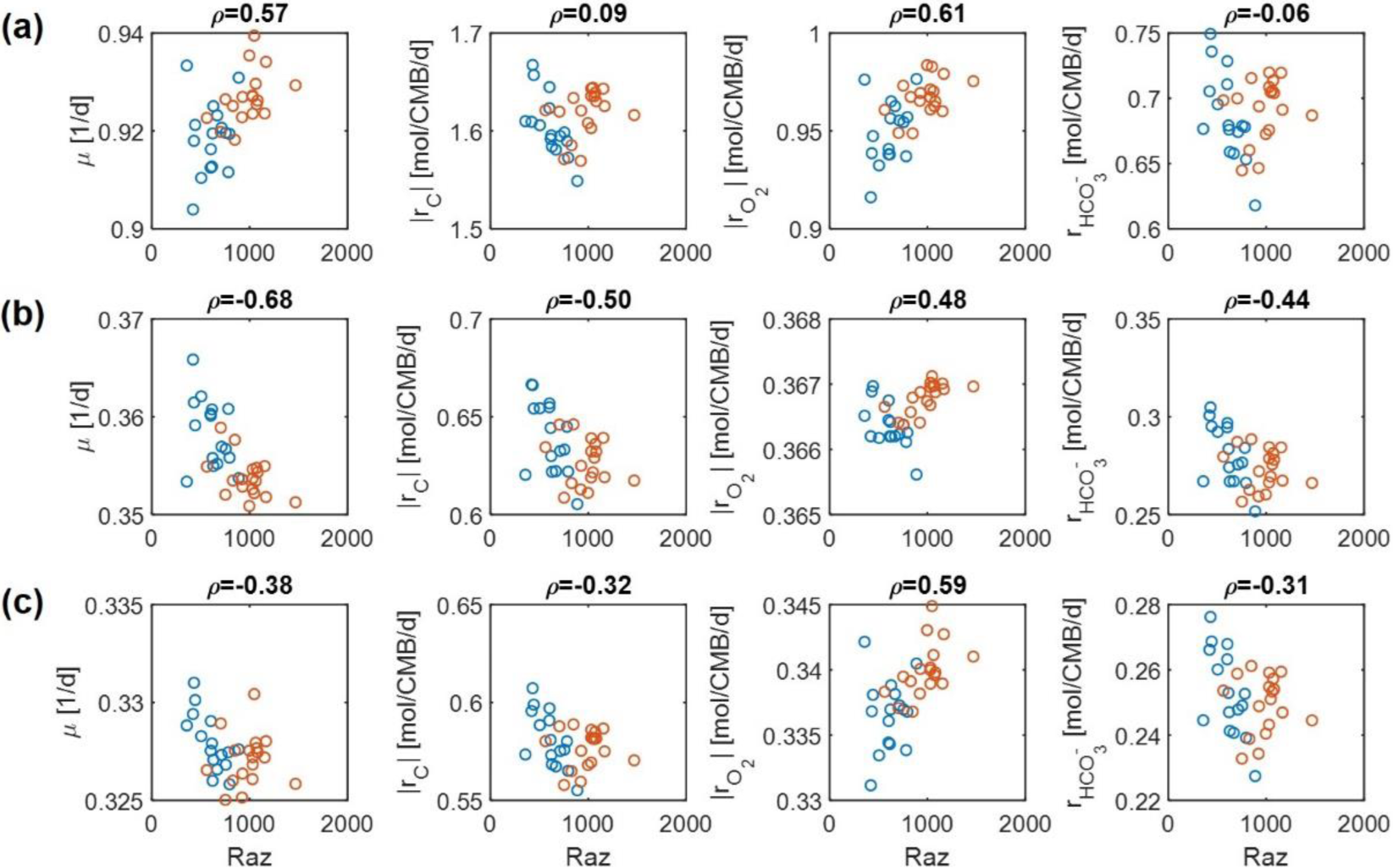
Pearson correlations of aerobic respiration with predicted reaction rates under moderate nutrient limitations (i.e., *V*_*h*_ [*OC*] = 1 and/or *V*_*h*_ [*O*_2_] = 1): growth rate (*μ*), carbon consumption rate (| *r*_*C*_ |), O_2_ consumption rate 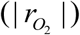, and bicarbonate production rate 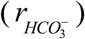: (a) C-limited condition, (b) O_2_-limited condition, and (c) both C&O_2_-limited condition.

**Figure 4.**
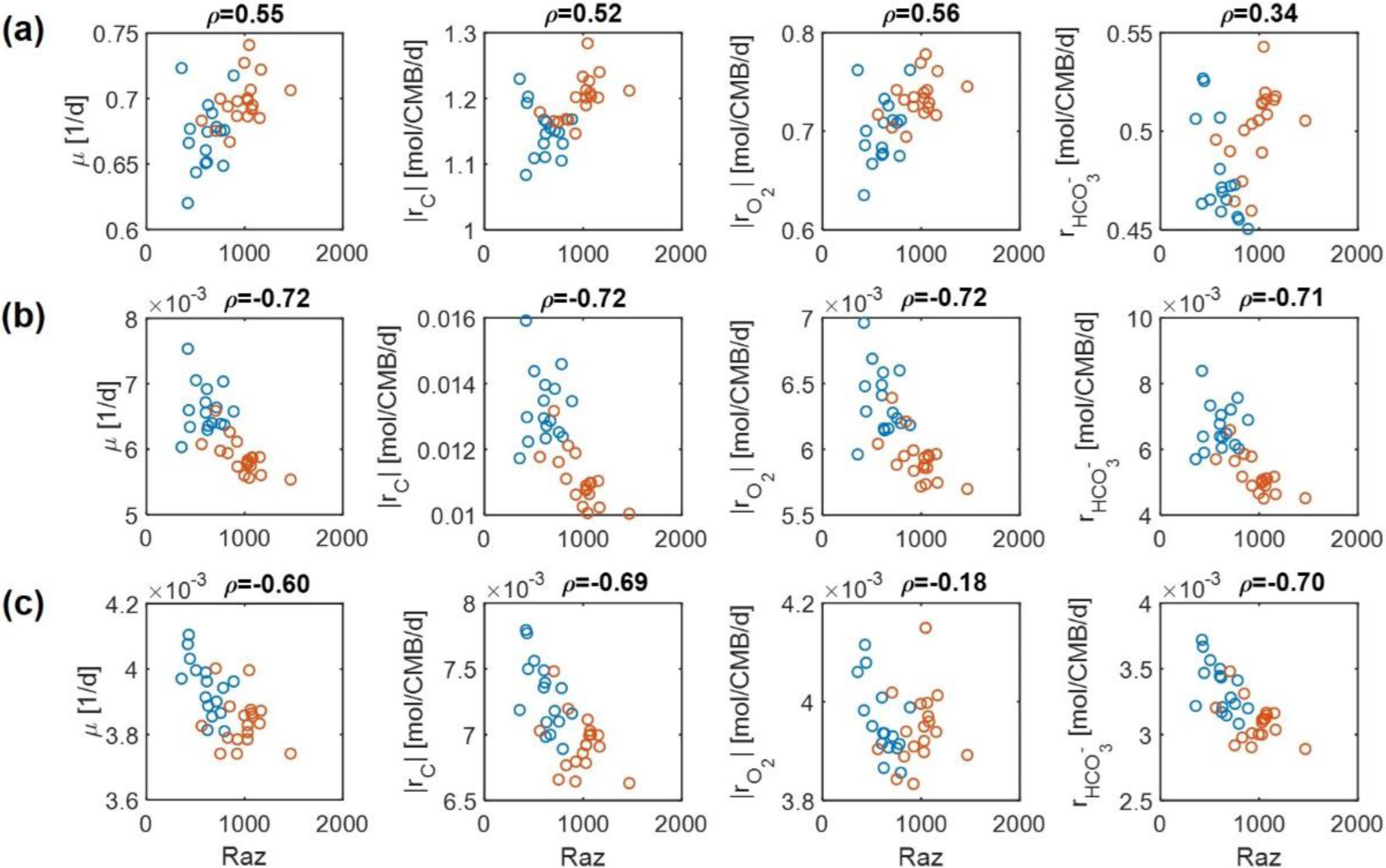
Pearson correlations of aerobic respiration with predicted reaction rates under severe nutrient limitations (i.e., *V*_*h*_ [*OC*] = 0.2 and/or *V*_*h*_ [*O*_2_] = 0.2): growth rate (*μ*), carbon consumption rate (| *r*_*C*_ |), O_2_ consumption rate 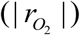, and bicarbonate production rate 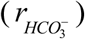: C-limited condition, (b) O_2_-limited condition, and (c) both C&O_2_-limited condition. For understanding the relationships between aerobic respiration and other rates, we analyzed multiple reaction rates as follows: biomass production (i.e., growth) rate (*μ*), C consumption rate (*r*_*C*_ ≡ (# of C) = *r*_*OC*_), O_2_ consumption rate 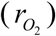, and inorganic carbon production rate 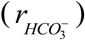 as defined in Eqs. (26) to (28).

For the case of moderate substrate limitation (Figure 3), for example, *μ* showed a positive correlation under the C-limited condition, which was however turned into negative correlations under O_2_- or both C&O_2_-limited conditions. Similar patterns were observed for | *r*_*C*_ | (i.e., the absolute value of *r*_*C*_) and 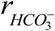, while their correlations with aerobic respiration were weak under the C-limited condition. As an exception, the correlation of 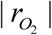 with aerobic respiration was consistently positive across all three different limiting conditions. As mentioned above, this result partially validates our model because Raz reduction to resorufin represents an estimate of the consumption rate of O_2_, rather than other chemicals (Gonzalez-Pinzon et al., 2012).

The results above were significantly changed when substrate limitation was severe (Figure 4). In this case, correlation patterns among four different reaction rates were shown to be similar. That is, they all showed positive correlations with aerobic respiration under C-limited conditions and negative correlations under O_2_- and both C&O_2_-limited conditions. Interestingly, the positive correlation of aerobic respiration under C-limited conditions was the highest for microbial growth and O_2_ consumption rates. Together with the previous result, this shows that experimental data in (Graham et al., 2017) may be consistently interpreted as being C-limited.

### Comparison of low-activity and high-activity samples

To understand how biogeochemistry in low- and high-activity zones is differentiated, we performed detailed analyses of two selected samples: one from the low-activity zone (sample N1-40-50) and the other from the high-activity zone (sample S1-00-10) (see Methods). With outliers removed, these two samples represent the highest and lowest activity as shown in the correlations of aerobic respiration with λ and 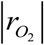 (Supplementary Figure S2).

Distributions of all three thermodynamic parameters (λ, Δ*G*^*Cat*^, and Δ*G*^*D*^) indicated that the OM pool at HA is more thermodynamically favorable than at LA (Figure 5). The distributions of parameter λ in the HA and LA samples, respectively, showed an exponential decay and bell-shaped patterns, indicating that the HA sample contained a predominantly large portion of thermodynamically favorable OM in comparison to the LA sample (Figure 5a). By contrast, the distributions of Δ*G*^*Cat*^(Figure 5b) and Δ*G*^*D*^(Figure 5c) suggested that the HA sample contained a lower portion of thermodynamically less favorable OM, compared to the LA sample.

**Figure 5.**
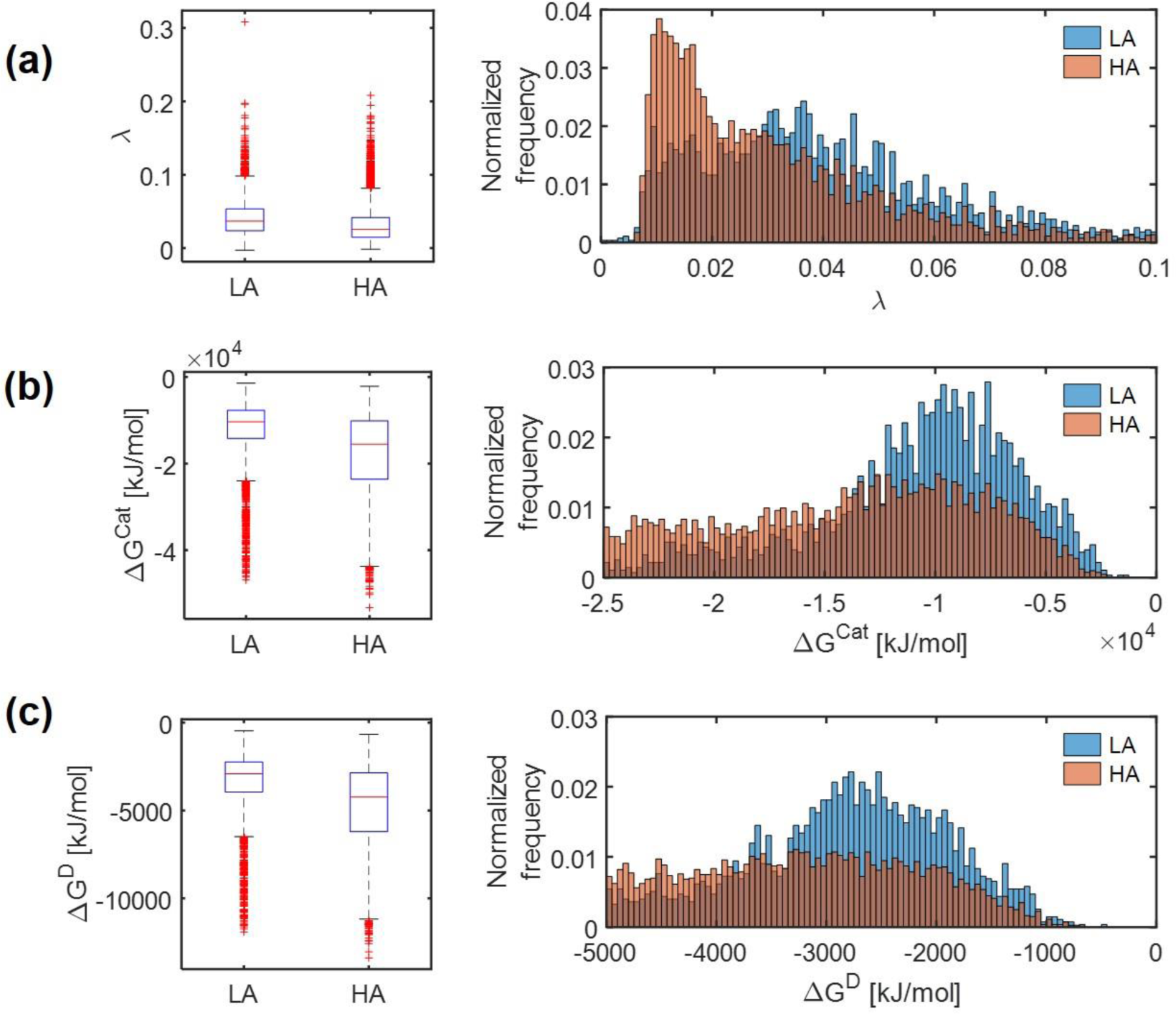
Box plots (left) and histograms for distributions of three key thermodynamic functions under: (a) Gibbs free energy change for an electron donor half reaction (Δ*G*^*D*^), (b) Gibbs free energy change for catabolic reaction (Δ*G*^*Cat*^), and (3) the energy coupling parameter (*λ*).

We further compared distributions of model-predicted oxidative respiration rates 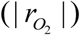 at LA vs. HA under C-, O_2_-, and both C&O_2_-limited conditions. In the case that substrates were *moderately* limited (i.e., *V*_*h*_ [*OC*] = 1 and/or *V*_*h*_ [*O*_2_] = 1) (Figure 6), predicted oxidative respiration at HA was higher than at LA, when C or both C&O_2_ were limited (as indicated by higher portions of faster 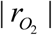 in the HA distribution) (Figures 6a and c); however, when O_2_ was limited, there was no significant difference between the two samples (Figure 6b).

**Figure 6.**
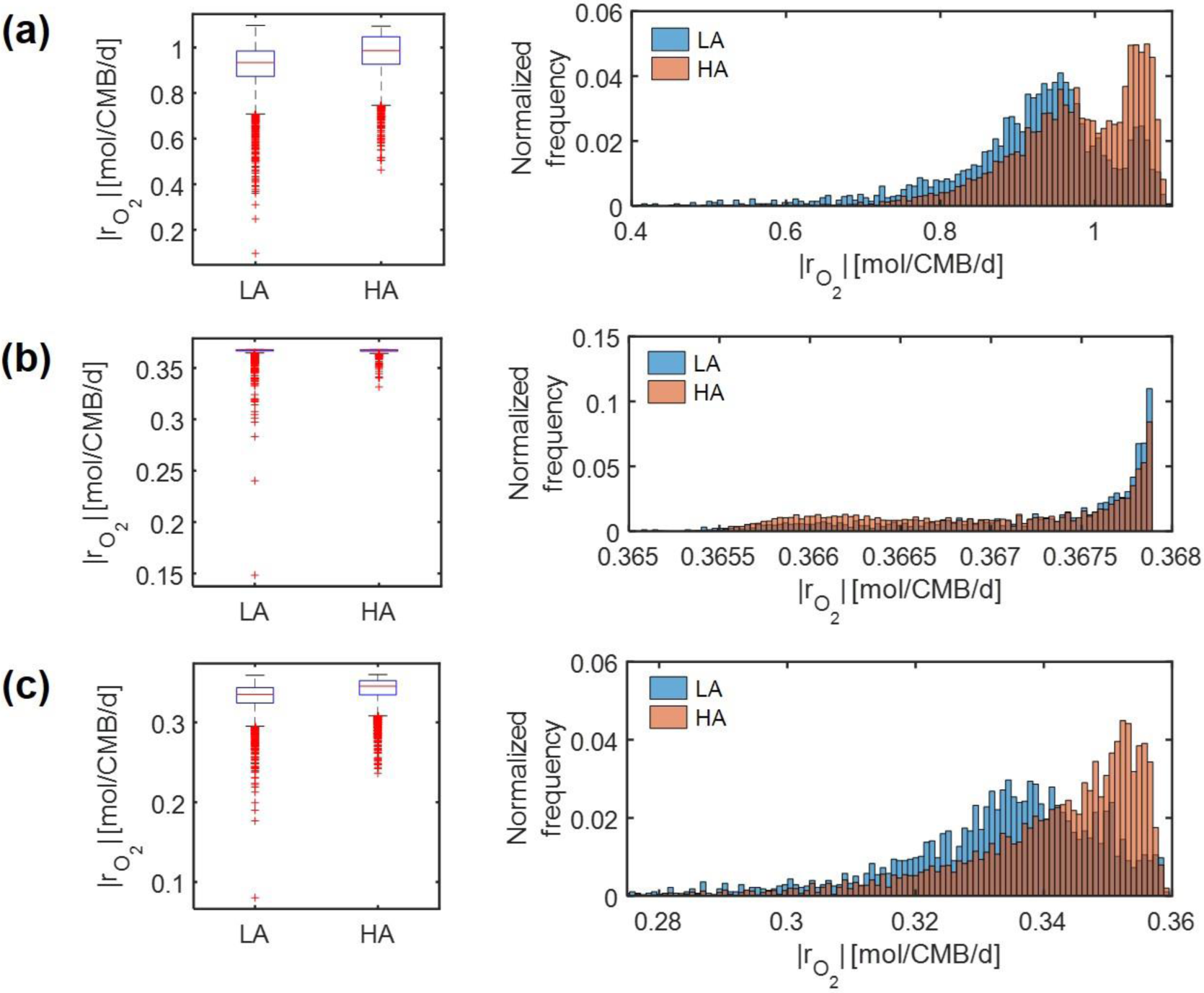
Box plots (left) and histograms for distributions of oxidative respiration rate 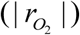 under moderate nutrient limitations (i.e., *V*_*h*_ [*OC*] = 1 and/or *V*_*h*_ [*O*_2_] = 1): (a) C-limited condition, O_2_-limited condition, and (c) both C&O_2_-limited condition.

Under *severe* substrate limitation (i.e., *V*_*h*_ [*OC*] = 0.2 and/or *V*_*h*_ [*O*_2_] = 0.2) (Figure 7), predicted oxidative respiration was higher at HA and at LA when C was limited (Figure 7a), while the increase in oxidative respiration at HA was moderate when O_2_ or C&O_2_ were limited (Figures 7a and 7b). Interestingly, when C was limited, 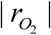 showed bimodal distributions in both HA and LA samples, indicating the nonlinear relationship between thermodynamic parameters to reaction rates.

**Figure 7.**
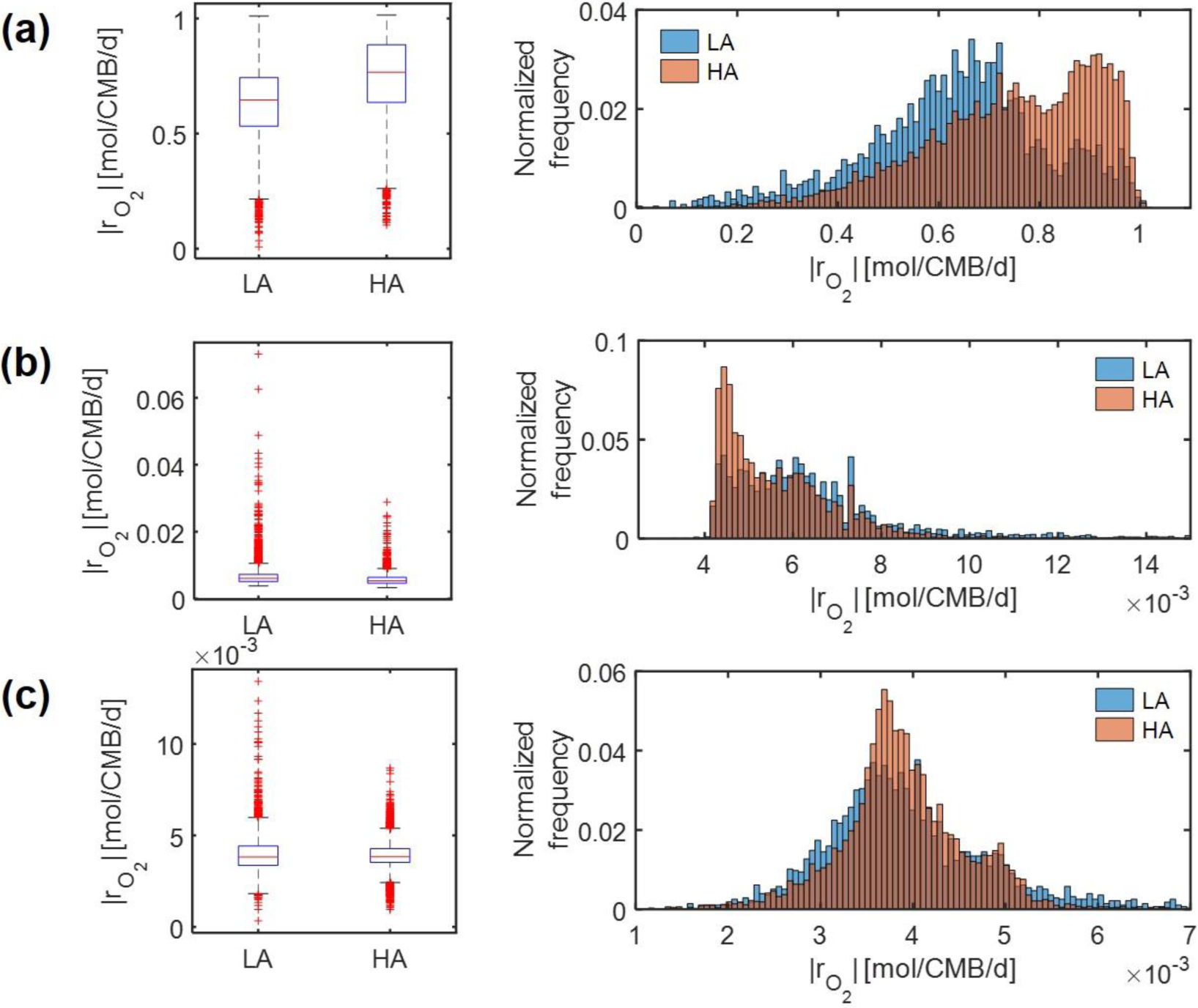
Box plots (left) and histograms for distributions of oxidative respiration rate 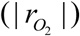 under severe nutrient limitations (i.e., *V*_*h*_ [*OC*] = 0.2 and/or *V*_*h*_ [*O*_2_] = 0.2): (a) C-limited condition, (b) O_2_-limited condition, and (c) both C&O_2_-limited condition.

Ratios of predicted reaction rates between the two samples (*HA* / *LA*) showed the level of elevated oxidative respiration in the HA zone (Figure 8). Although these ratios varied among reaction rates and were affected by the level of substrate limitation, their trends across three different substrate-limited conditions were consistent. That is, in cases of moderate (Figure 8a) and severe (Figure 8b) substrate limitation, we consistently found: (1) that the ratios of reaction rates were lowest under O_2_ limitation, intermediate when both C and O_2_ were limited, and highest under C limitation; and (2) that the magnitude of the ratio always maintained the following order: 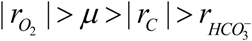.

**Figure 8.**
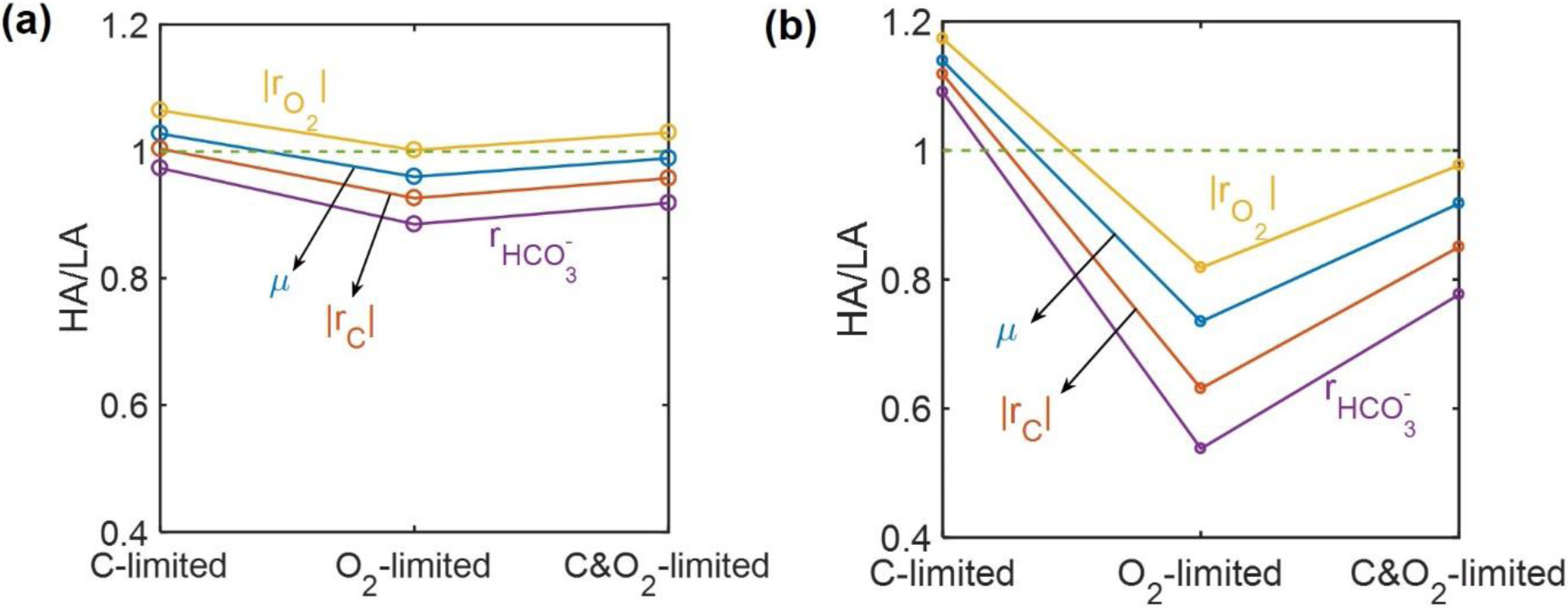
Reaction ratios between the HA and LA samples under C-, O_2_-, and C&O_2_-limited conditions: (a) moderately limited (i.e., *V*_*h*_ [*OC*] = 1 and/or *V*_*h*_ [*O*_2_] = 1), and (b) severely limited (i.e., *V*_*h*_ [*OC*] = 0.2 and/or *V*_*h*_ [*O*_2_] = 0.2).

Lastly, we performed dynamic simulations of OM consumption in LA and HA using two types of models: (1) SXM, and (2) a hybrid model that integrates SXM and EXM. The latter is a special case of a general IBM platform that hybridizes three complementary modeling platforms approaches: SXM-EXM-MXM. In hybridizing SXM with EXM, we used the cybernetic approach (Ramkrishna and Song, 2012;Ramkrishna and Song, 2018) to account for regulation of enzyme synthesis to simulate selective activation of oxidative reactions among a number of pathways (Methods), a key aspect that was missing in the case of using the SXM alone. Although both SXM and hybrid models correctly predicted that specific oxidative reaction rates at HA were higher than at LA (Supplementary Figure S3), the time scale of the SXM was shown to be very small, which led specific reaction rates to be unrealistically high (i.e., more than 300 mol/CMB/d) (Supplementary Figures S3a and S3b). By contrast, the hybrid model predicted specific reaction rates consistent with the literature values (Supplementary Figures S3c and S3d). We further compared the two models with respect to how specific OM consumption rates would change with the number of incorporated compounds. As *specific* rates are defined per unit C-mole of biomass, their dependency on the number of compounds is expected to be insignificant, but specific rates calculated by the SXM linearly increased with the number of compounds (Figure 9a), while the hybrid model predicted specific rates to be almost constant regardless of the number of compounds (Figure 9b). These results together indicate that SXM-EXM coupling significantly improved model predictions.

**Figure 9.**
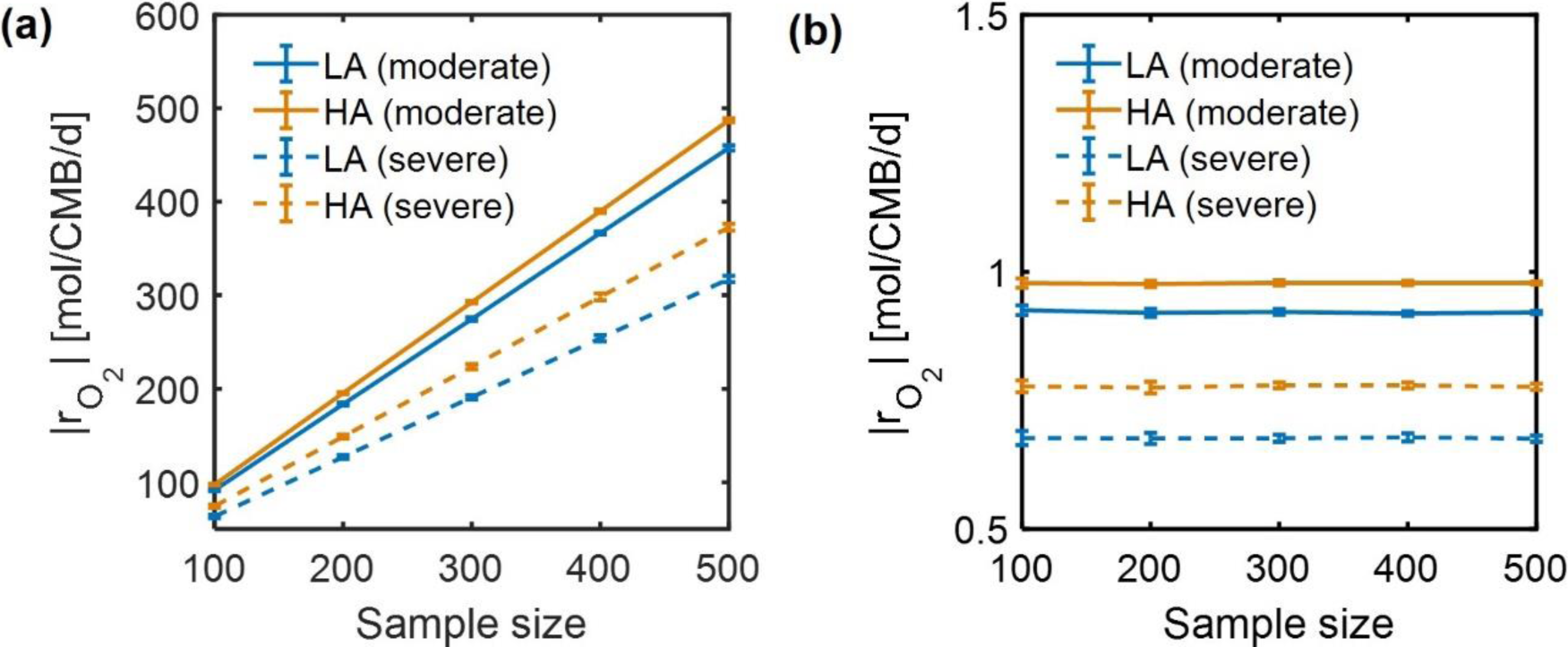
Dependence of simulated respiration rates 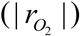 using: (a) SXM and (b) hybrid model (SXM+EXM).

## DISCUSSION

In this work, we proposed a novel and flexible biogeochemical modeling concept, termed substrate-explicit modeling (SXM), that accounts for the chemistry of individual OM molecules by using thermodynamic properties to distill this complexity into two essential parameters. A key feature of SXM is its ability to incorporate data from increasingly high-resolution metabolomics technologies into biogeochemical models by formulating OM-specific reaction kinetics for an unlimited number of organic compounds in a sample. The entire set of the resulting reaction kinetics is then represented by only two parameters (maximal growth rate and harvest volume). Our framework is a unique and scalable tool for modeling complex biogeochemical cycles at the ecosystem-scale, as no other approach can describe dynamic biogeochemical reaction networks composed of thousands of compounds with a small, computationally feasible set of parameters.

In application to field data, we showed aerobic respiration as being driven by thermodynamic favorability of compounds. This result challenges classical theories that use the concentrations of bulk substrate pools (such as organic carbon and oxygen) as the sole driving factors of aerobic respiration, notably excluding the influence of OM chemistry on biogeochemistry. The substrate-explicit model is built upon recent experimental studies that reveal a close relationship between OM thermodynamics and aerobic respiration (Graham et al., 2017;Graham et al., 2018;Garayburu-Caruso et al., 2020). Our test cases are broadly consistent with the conclusions of these studies, indicating a need to revise classic theory via incorporation of OM thermodynamics into our understanding and modeling of aerobic respiration.

We therefore suggest the use of λ as a new thermodynamic parameter to evalute thermodynamic character of OM, and model the biogeochemical implications of variation in those thermodynamics. It has been customary to use Gibb free energy for an electron donor half reaction (Δ*G*^*D*^) at a standard condition (pH=0) (often denoted by 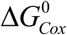 in the literature) (LaRowe and Van Cappellen, 2011). This does not, however, properly represent conditions where actual biochemical reactions occur and thus needs be corrected (e.g., to pH=7). Surprisingly, the correlations of aerobic respiration with Δ*G*^*D*^ for pH=0 and pH=7 showed fundamentally different trends, among which the latter (i.e., at pH=7) provided the more interpretable results. In contrast with Δ*G*^*D*^, which considers energy generation only through an electron donor half reaction, λ accounts for the energy balance between catabolic and anabolic reactions, thus evaluating thermodynamic properites of compounds based on the complete chemistry. Indeed, λ consistenly showed negative correlations with aerobic respiration (meaning that thermodynamically more favorable OM shows higher respiration rate), regardless of pH values.

Our model was validated through a consistency check between predicted oxygen consumption rates with experimentally determined aerobic respiration. Across all three conditions (C, O_2_ and, both C&O_2_ limitation), aerobic respiration showed posistive correlations with oxygen consumption rate when substrates are moderately limited. By contrast, in the case of severe C and O limitation, a positive correlation bewteen oxygen consumption rate and aerobic respiration was obtained only for C-limited conditions. This matches field conditions for the study system in which OC concentrations are low and porewater within saturated sediments is consistently aerobic (Stegen et al., 2018b). These results together suggest that the field data is consistently interpretable due to C limitation and O_2_ excess. In support of this, recent experimental work has shown a dependency of aerobic respiration on OM thermodynamics when C is limited but has shown no effect of C thermodynamics on respiration when C is widely available (Garayburu-Caruso et al., 2020).

The difficulty in finding reliable kinetic parameters is often a major hurdle in scaling up biogeochemical models to large-scale complex systems. As mentioned earlier, this barrier is overcome by our modeling approach that formulates compound-specific reaction rates with only two parameters (maximal growth rate and harvest volume). While accurate determination of those two parameters is ideally pursued via experimental data, values determined by a thermodynamic transition theory can serve as a reasonable proxy. As such, our model can guide experimental design to refine parameters and understand factors driving their variation across systems.

Lack of quantitative information (i.e., concentrations) on individual OM molecules is an intrinsic limitation in analyzing OM profiles from FTICR-MS. Consequently, we focused on evaluating carbon *quality* based on the normalized distribution of OM, but this gap can be be filled for improving predictions, e.g., through the integration with other complementary metabolomics approaches that can provide quantitative data (Hertkorn et al., 2013;McCallister et al., 2018).

The capability of our method that incorporates high-resolution mass spectrometry (or OM characterization methods) data into biogeochemical modeling greatly facilitates other research programs in the field that collect OM chemistry datasets. The Worldwide Hydrobiogeochemistry Observation Network for Dynamic River Systems (WHONDRS), for example, is a global research consortium that aims at understanding cross-scale dynamic interactions among hydrology, biogeochemistry, and microbiology in river corridors (Stegen and Goldman, 2018). As an initial effort, WHONDRS provides the collection of high-resolution OM profiles such as FTICR-MS data across rivers in the world (Stegen et al., 2018a;Chu et al., 2019;Danczak et al., 2019;Garayburu-Caruso et al., 2019;Goldman et al., 2019;Renteria et al., 2019;Stegen et al., 2019;Wells et al., 2019;Danczak et al., 2020). Other networked efforts, such as the National Ecological Observation Network (Teeri and Raven, 2002;Barnett et al., 2019), provides similar kinds of data that are amenable for analysis vis SXM. Analysis of these data, which are collected and analyzed consistently across systems, using the SXM framework will significantly improve our understanding of the level of heterogeneity across space and time in OM consumption and respiration, and thus could be used as a critical tool for more mechanistic predictions of spatial and temporal variation in stream/river CO_2_ emissions and other coupled biogeochemical rates from local to global scales.

As shown, a synergistic integration of SXMs with other existing frameworks, such as MXMs and EXMs, is an important future direction to fill the gaps that each approach has and to address advanced science questions that could not be addressed individually. We illustrate a conceptual master platform of integrative biogeochemical modeling (IBM) that accounts for the interaction among substrates, microbes, and enzymes at an unprecedented level of detail (Figure 10). We envision that the IBM will significantly contribute to create a molecular-level understanding of biogeochemistry that can be translated to complex ecosystem modeling. In this regard, the SXM concept propsed in this work provides a key component of IBM that has been missing so far.

**Figure 10.**
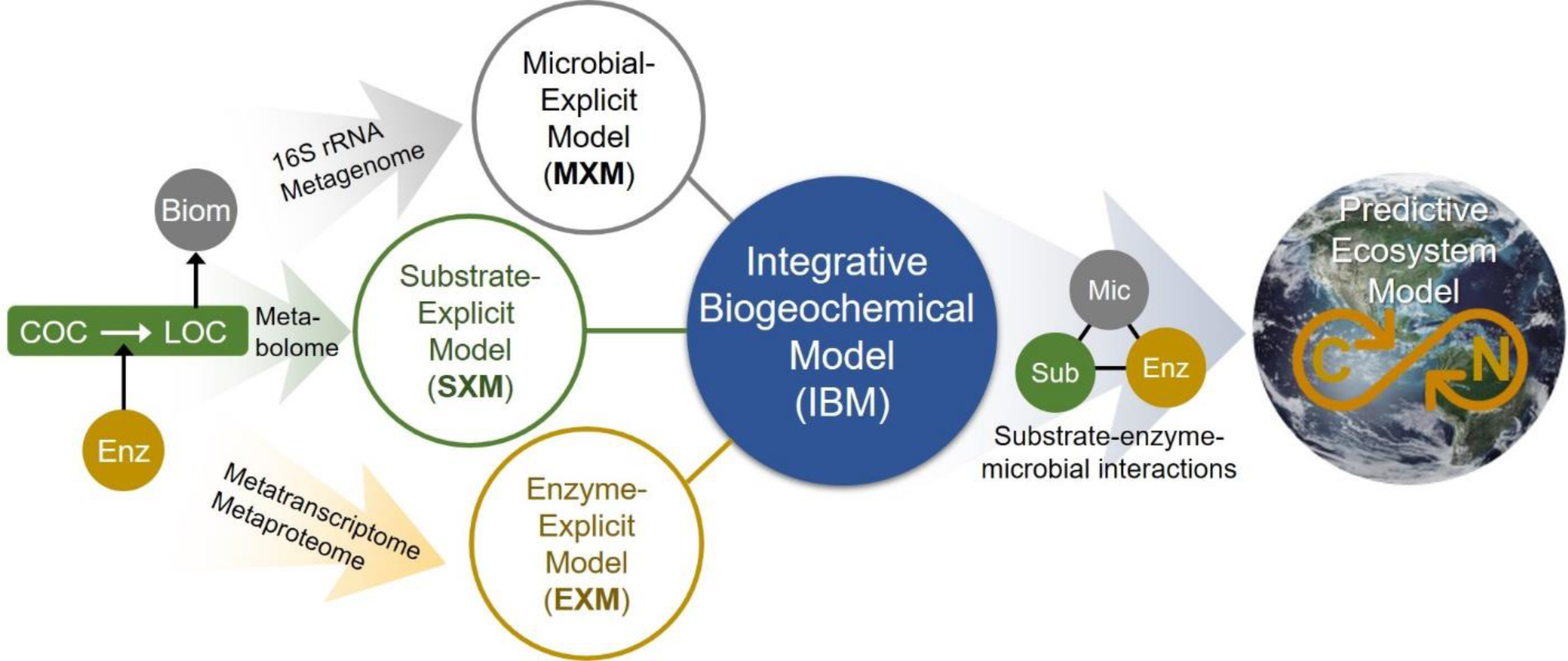
The integrated biogeochemical modeling (IBM) concept that combines MXM, SXM, and EXM, which are respective representations of significant expansions from typical lumped models by integrating multi-omics data to identify functional contributions of individual organisms and/or functional guilds, substrate-specific degradation pathways, and detailed enzymatic processes. Consequently, the IBM may enable providing a mechanistic understanding of dynamic linkage and interactions among substrates, enzymes, and microbes at a molecular level and significantly improving the performance of complex ecosystem models in predicting OC consumption and CO_2_ emission in space and time.

## Author contributions

H.-S.S., J.C.S., E.B.G., V.G.-C., and T.D.S. conceptualized the study; H.-S.S. developed the codes and performed theoretical analyses. J.-Y.L., and W.C.N. prepared data files; H.-S.S. drafted out the manuscript, which was edited by J.C.S and E.B.G.; All authors contributed to the writing.

## Acknowledgments

This research was supported by the U.S. Department of Energy (DOE), Office of Biological and Environmental Research (BER), as part of Subsurface Biogeochemical Research Program’s Scientific Focus Area (SFA) at Pacific Northwest National Laboratory (PNNL). PNNL is operated for DOE by Battelle under contract DE-AC06-76RLO 1830. A portion of the research was performed at the Environmental Molecular Science Laboratory User Facility located on PNNL’s campus.

---

**Supp Figure S1.**
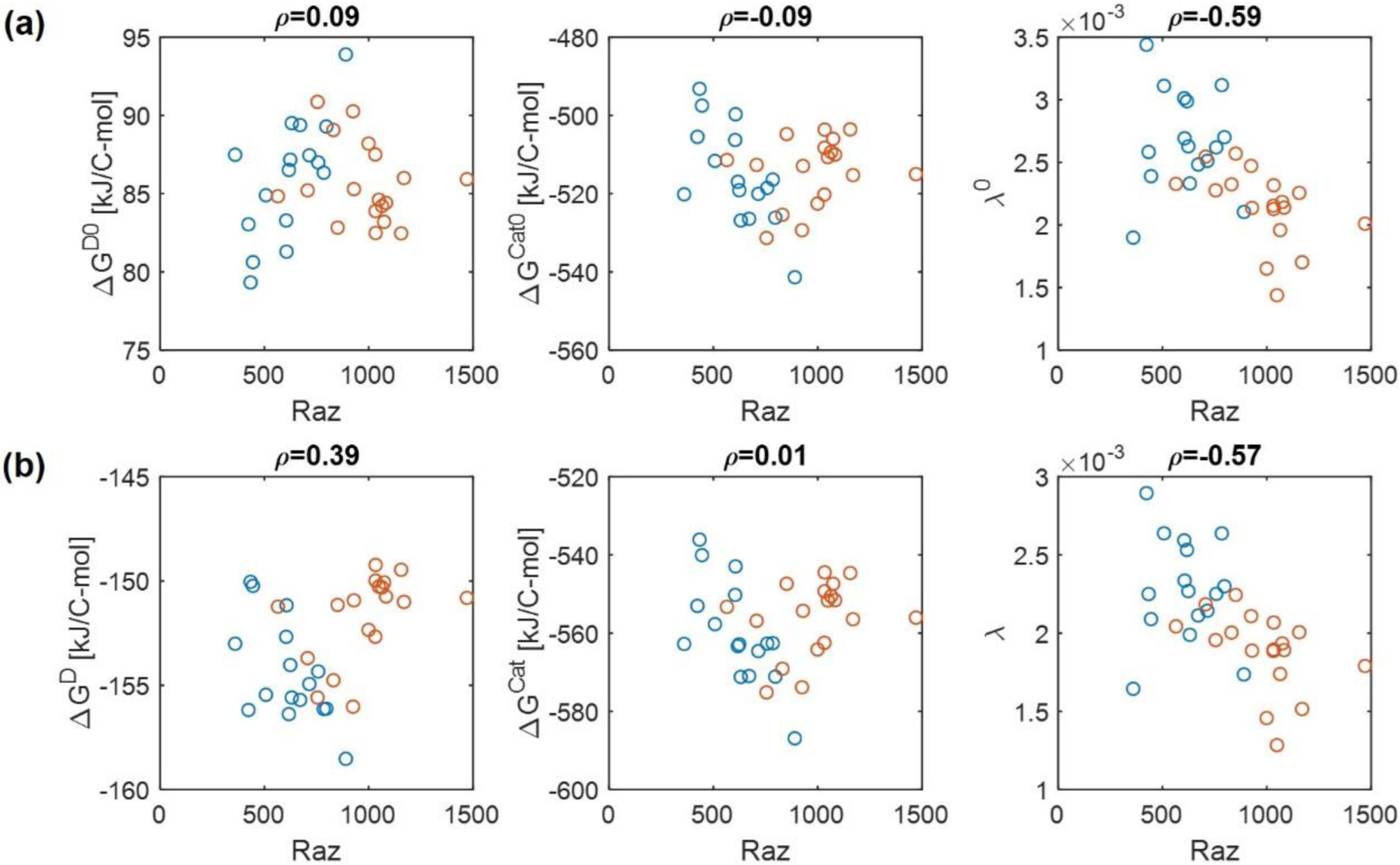
Replot of Figure 2 with recalculation of thermodynamic functions per a unit C-mole of OC: (a) pH=0 (denoted by the superscript 0) and (b) pH=7.

**Supp Figure S2.**
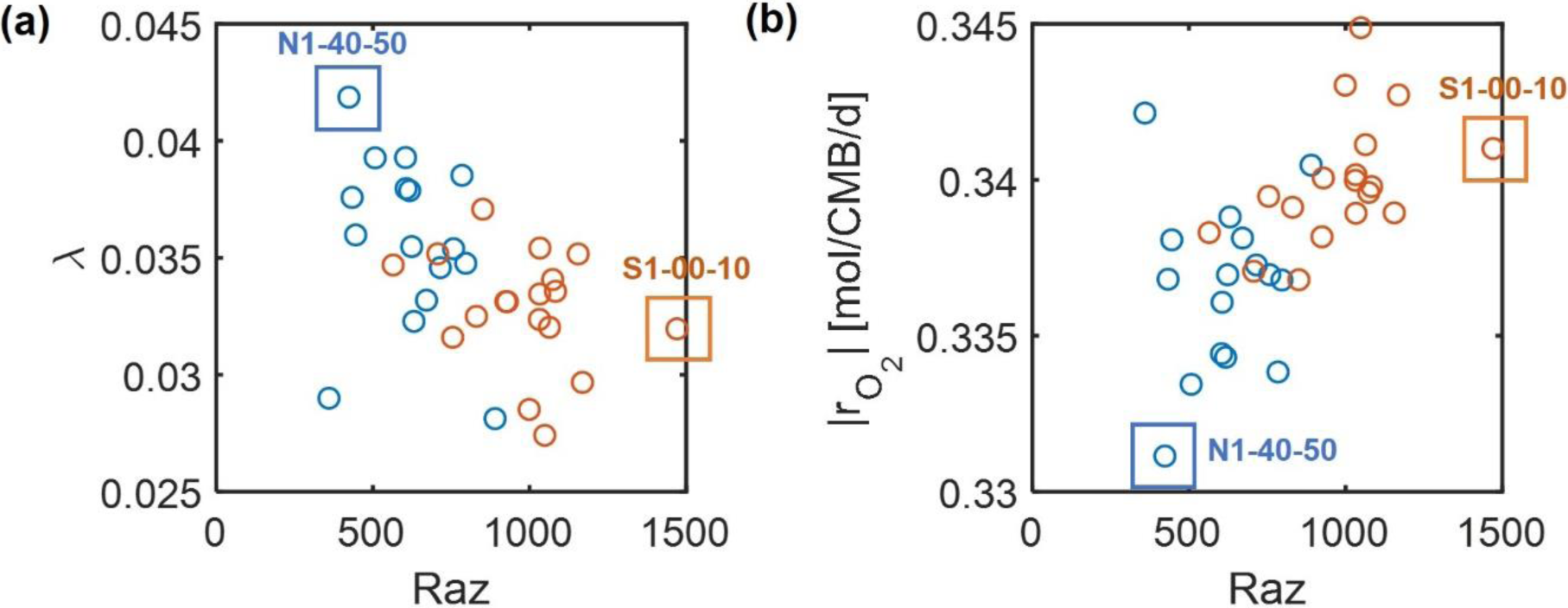
Choice of two representative sections for the comparison of the low- and high-activity zones based on the correlations of aerobic respiration with (a) the energy coupling parameter (*λ*) and O_2_ consumption rate 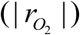.

**Supp Figure S3.**
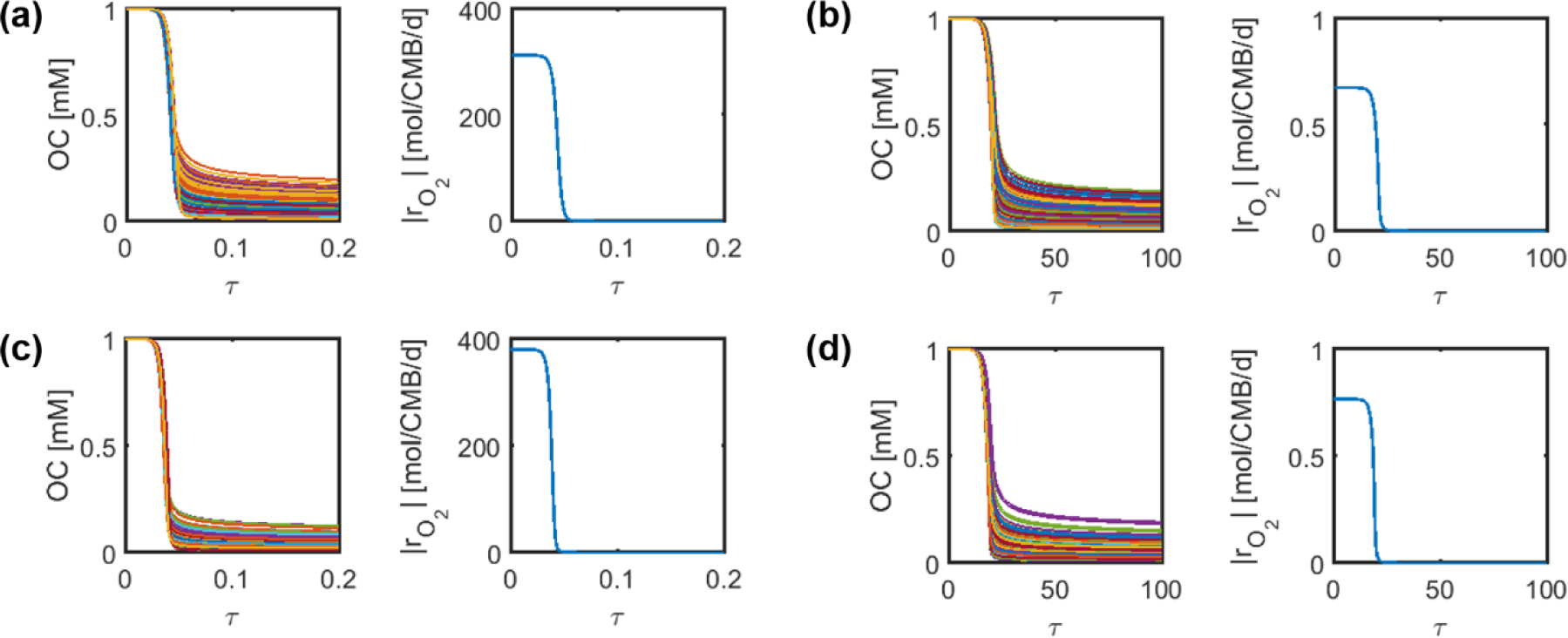
Dynamic simulations based on 500 OC randomly selected respectively from the LA and HA nutrient pools: (a) SXM for the LA sample, (b) hybrid model (SXM+EXM) for the LA sample, (c) SXM for the HA sample, (d) hybrid model (SXM+EXM) for the HA sample. *τ* denotes the dimensionless time (≡ *tμ*^max^).

